# Radiation-Induced Changes in Energy Metabolism Result in Mitochondrial Dysfunction in Salivary Glands

**DOI:** 10.1101/2023.11.27.568879

**Authors:** Lauren G. Buss, Brenna A. Rheinheimer, Kirsten H. Limesand

**Affiliations:** School of Nutritional Sciences and Wellness, University of Arizona, Tucson, AZ, United States of America; University of Arizona Cancer Center, Tucson, AZ, United States of America

## Abstract

Salivary glands are indirectly damaged during radiotherapy for head and neck cancer, resulting in acute and chronic hyposalivation. Current treatments for radiation-induced hyposalivation do not permanently restore function to the gland; therefore, more mechanistic understanding of the damage response is needed to identify therapeutic targets for lasting restoration. Energy metabolism reprogramming has been observed in cancer and wound healing models to provide necessary fuel for cell proliferation; however, there is limited understanding of alterations in energy metabolism reprogramming in tissues that fail to heal. We measured extracellular acidification and oxygen consumption rates, assessed mitochondrial DNA copy number, and tested fuel dependency of irradiated primary salivary acinar cells. Radiation treatment leads to increases in glycolytic flux, oxidative phosphorylation, and ATP production rate at acute and intermediate time points. In contrast, at chronic radiation time points there is a significant decrease in glycolytic flux, oxidative phosphorylation, and ATP production rate. Irradiated salivary glands exhibit significant decreases in spare respiratory capacity and increases in mitochondrial DNA copy number at days 5 and 30 post-treatment, suggesting a mitochondrial dysfunction phenotype. These results elucidate kinetic changes in energy metabolism reprogramming of irradiated salivary glands that may underscore the chronic loss of function phenotype.

## INTRODUCTION

According to the National Cancer Institute’s United States (US) Cancer Statistics data, the prevalence of oropharyngeal cancer has increased over the past 20 years in the US with the highest increases observed in older males ^1,2^. With the prevalence rate increasing, the quality of life of head and neck cancer (HNC) patients is becoming increasingly important to address as the number of cancer survivors continues to grow due to improvements in diagnosis and treatment ^3,4^. Human papillomavirus (HPV)-associated HNC cases have increased as smoking-associated HNC cases have decreased, and HPV-associated HNC patients display significantly increased progression-free survival compared to smoking-associated HNC cancer patients ^5,4,6^. Although HPV-associated HNC patients exhibit increased long-term survival, they still experience adverse side effects associated with radiation therapy (IR), which is the standard of care for approximately 80% of HNC patients ^7,8^. The salivary gland is a nearby organ in HNC patients that is damaged by IR, resulting in hyposalivation, mucositis, xerostomia, malnutrition, and dental caries ^9,10,11^. The majority of HNC patients exhibit xerostomia (70-80%) that lasts chronically, which negatively affects their quality of life ^9,10,11^. Xerostomia is typically managed with topical saliva substitutes or cholinergic drugs that stimulate saliva production such as pilocarpine and cevimeline, but they only temporarily manage the symptoms and cholinergic agents have been reported to produce unpleasant side effects such as sweating and gastrointestinal distress ^7,10,12^. New treatments are needed to restore function to salivary glands in order to treat xerostomia in HNC patients, therefore more mechanistic understanding of radiation-induced salivary gland damage is needed to identify therapeutic drug targets to restore tissue function.

Energy metabolism is a well-studied mechanism that is necessary to produce ATP, amino acids, and fatty acid intermediates to fuel biosynthetic processes within cells to restore damaged tissue. The association between aberrant energy metabolism and disease has been established in type II diabetes mellitus, cancer, aging, and organ fibrosis ^13,14,15,16^. In the wound-healing response, glycolytic activity is observed to increase quickly to synthesize ATP and generate carbon backbones for macromolecule synthesis in stem cells, muscle tendons, endothelial cells, and epithelial cells ^17,18,19,20^. We previously integrated transcriptomic and metabolomic analysis and identified enrichment of energy metabolism at day 5 post-radiation treatment in the parotid salivary glands of mice ^21^, however, the effect of IR on glycolysis in the salivary gland has not been measured, which may uncover a new mechanism for restoring tissue function. Following IR treatment, acinar cells in the parotid salivary gland increase compensatory proliferation starting at day 5 and this increase is sustained chronically ^22,23,24^. Increased compensatory proliferation correlates with loss of secretory function ^24^; therefore, it is essential to investigate if glycolysis is a mechanism driving this sustained compensatory proliferation.

Mitochondria are critical organelles in metabolic processes both under homeostasis and repair conditions. It has been established that reactive oxygen species (ROS) levels increase acutely in the mitochondria of irradiated salivary gland acinar cells ^25^ and targeting this source of ROS using MitoTEMPO prior to radiation treatment preserves salivary gland function ^26^.

Improper electron transport chain function has been reported as an indicator of mitochondrial dysfunction ^27,28,29^. In our previous work, we identified reductions in NAD+, UQCR, NDUF, COX, ATP5, and MT-ND family members in the salivary gland five days following targeted 5 Gy IR, suggesting defects in the function of complexes I, III, IV, and V within the electron transport chain ^21^. Additionally, an increase in mtDNA copy number (polyploidization) is believed to be an adaptive repair response when mitochondrial function is reduced due to ROS damage ^30^. Although mtDNA copy number polyploidization has been observed following IR, the effects of this process on mitochondrial oxidative phosphorylation are not understood.

The objective of this study was to investigate changes in energy metabolism and mitochondrial function over time in the parotid salivary glands of mice following a single 5 Gy dose. This radiation dose mimics the clinical fraction of 2 Gy/day, and we hypothesize that metabolic changes observed early in the treatment regimen set the environment for inefficient repair and chronic loss of function. We demonstrate that energy metabolism increases early (acute and intermediate time points) in primary salivary acinar cells harvested from mice, which is attenuated at chronic time points. Radiation-induced decreases in spare respiratory capacity and increases in mtDNA copy number at intermediate or chronic time points underscore the continual decline in mitochondrial function. These data suggest that chronic energy metabolism impairment and mitochondrial dysfunction are important phenotypes involved in radiation-induced loss of tissue function.

## RESULTS

### Cellular bioenergetics of irradiated salivary acinar cells increase at acute time points

Measurements of the extracellular acidification rate (ECAR) and oxygen consumption rate (OCR) of live cells qualitatively reflects glycolysis and mitochondrial respiration, respectively ^31,32,33^. To understand the effect of radiation on cellular energy production in the salivary gland, the Seahorse XF Real-Time ATP Rate Assay Kit ^34^ was performed to collect ECAR and OCR data in salivary gland primary acinar cells and calculate basal ATP production rates at acute (24 and 48 hours) radiation (IR) time points (Fig. 1a for experimental design).

**Figure 1.**
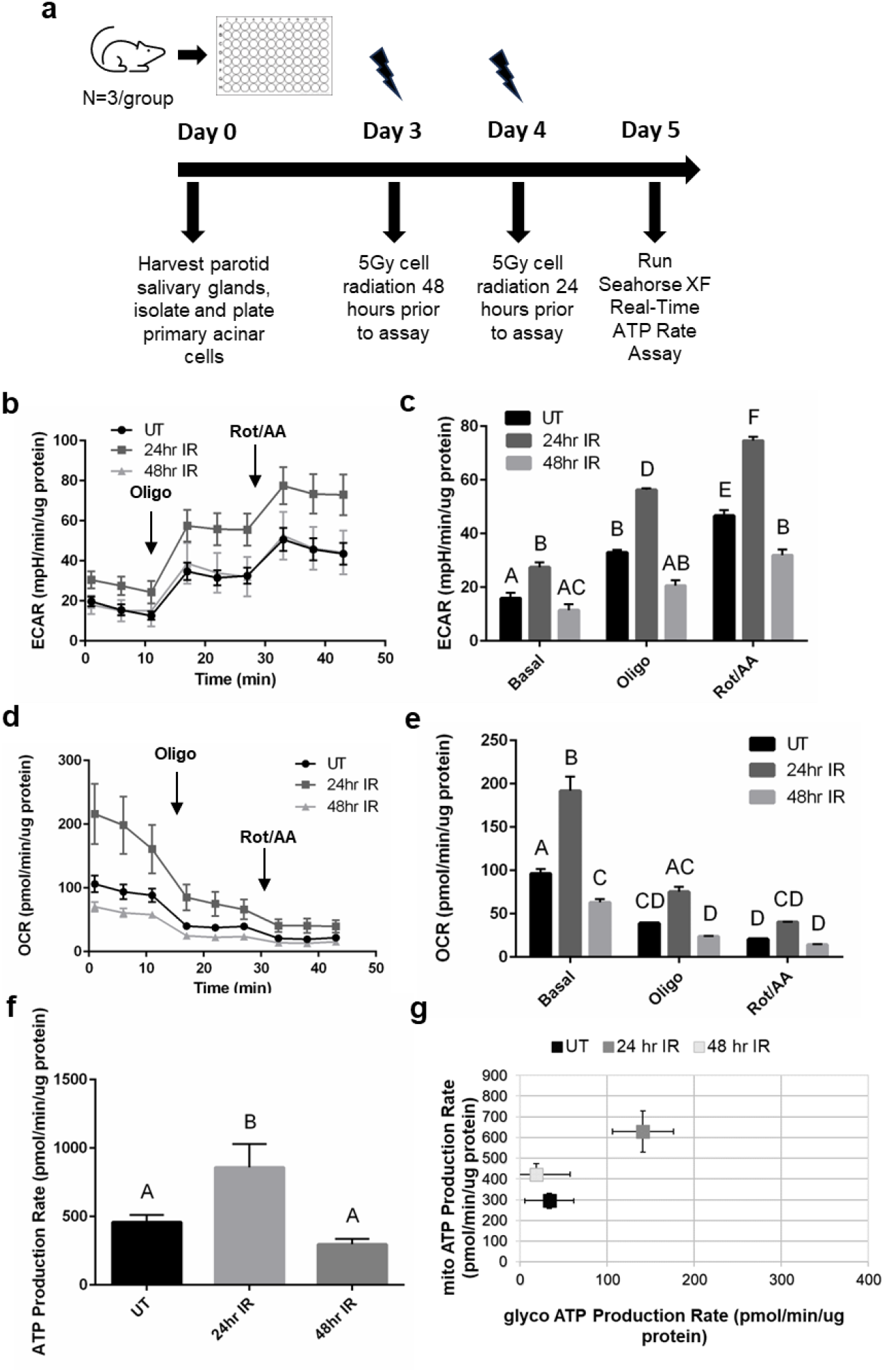
ECAR, OCR, and ATP production rate increase acutely at 24-hours following radiation in salivary acinar cells. (**a**) Primary acinar cells were isolated from parotid salivary glands of untreated female FVB mice (n=3/group), cultured for 5 days, and then irradiated at 24 and 48 hours prior to the Seahorse XF Real-Time ATP Rate Assay. (**b**) Individual ECAR readings for untreated (UT), 24-hour IR (24hr IR), and 48-hour IR (48hr IR) groups with the responses to 2.0uM oligomycin (oligo) injection and 0.5uM rotenone/antimycin A (rot/AA) injection shown. (**c**) Average basal, oligo, and rot/AA ECAR readings for UT, 24hr IR, and 48hr IR. (**d**) Individual OCR readings for UT, 24hr IR, and 48hr IR groups with the responses to 2.0uM oligomycin (oligo) injection and 0.5uM rotenone/antimycin A (rot/AA) injection shown. (**e**) Average basal, oligo, and rot/AA OCR readings for UT, 24hr IR, and 48hr IR. (**f**) Basal ATP production rates of UT, 24hr IR, and 48hr IR groups. (**g**) The proportion of mitochondrial (mito) ATP production rate and glycolytic (glyco) ATP production rate for UT, 24hr IR, and 48hr IR groups. Each panel is representative of 3 independent assays (n=3/group). Significant differences were determined using one-way ANOVA and Bonferroni post-hoc test, p<0.05. Treatment groups with the same letter are not significantly different from each other.

Primary cells were utilized due to the role of p53 signaling in the radiation response of salivary glands ^35,36^ and its potential role in regulating metabolic processes ^37^. Within irradiated salivary glands, apoptosis of acinar cells occurs 8-72 hours post-treatment with a peak occurring at 24 hours, and increased levels of ROS are observed at 1 and 3 days post-IR ^35,38,25^, therefore 24 and 48 hours post-IR were chosen as acute time points in this study. In this assay, oligomycin was injected into the cell media to inhibit mitochondrial ATP production and calculate the ATP production rate attributed to mitochondrial oxidative phosphorylation production while rotenone plus antimycin A was injected to completely inhibit mitochondrial respiration and is used with ECAR to calculate the ATP production rate attributed to glycolysis ^34^. We determined that the contribution of non-glycolytic acidification to ECAR is minimal by adding 2-deoxyglucose (2- DG) to the cells in an optimization experiment (Supplemental Fig. S1; for all supplemental material, use the link provided in the Supplemental Data section of the Methods). These data allow us to use the ECAR measurements from the ATP Rate Assay to properly assess glycolytic acidification^31,39^.

At baseline readings, 24-hour IR ECAR measurements are significantly higher compared to untreated and 48-hour IR (Fig. 1b,c). Following oligomycin and rotenone/antimycin A injections, 24-hour IR ECAR measurements remain significantly higher compared to untreated and 48-hour IR (Fig. 1b,c). At baseline readings, 24-hour IR OCR measurements are significantly higher than untreated and 48-hour IR OCR measurements are significantly lower compared to untreated (Fig. 1d,e). Following oligomycin and rotenone/antimycin A injections, the 24-hour IR OCR measurements remain higher compared to the 48-hour IR and untreated OCR measurements and the 48-hour IR and untreated OCR readings are not significantly different from each other (Fig. 1d,e). Calculation of the basal ATP production rates reveals a significantly higher ATP production rate in the 24-hour IR group compared to untreated and 48- hour IR groups (Fig. 1f), with the 24-hour IR group displaying higher glycolytic and mitochondrial ATP production rates compared to both the untreated and 48-hour IR groups (Fig. 1g). Acutely, we observe an increase in ECAR, OCR, and ATP production rates at 24 hours post-IR that is attenuated at 48 hours post-IR.

### Cellular bioenergetics of irradiated salivary acinar cells increase at an intermediate time point and decrease at chronic time points

Due to the observed changes in ECAR, OCR, and basal ATP production at acute time points following IR, we further investigated acinar cell bioenergetics at days 5, 30, and 60 following IR (Fig. 2a for experimental design). Day 5 post-IR was chosen as an intermediate time point where increased compensatory proliferation and decreased apical/basolateral polarity in the acinar compartment of the salivary gland begin following loss of function starting at day 3 post-IR ^23,40,41,42^. We have previously observed chronic loss of salivary gland function correlated with decreased acinar cell differentiation markers and continued compensatory proliferation at days 30, 60, and 90 post-IR ^43,40,23^, therefore days 30 and 60 were chosen as representative chronic IR time points. At baseline readings, day 5 IR ECAR measurements are marginally higher compared to untreated (Fig. 2b,c), and following rotenone/antimycin A injection the increase becomes statistically significant (Fig. 2b,c). At baseline readings, day 5 IR OCR measurements are significantly higher than untreated (Fig. 2d,e). Following oligomycin and rotenone/antimycin A injections, day 5 IR OCR measurements remain higher compared to untreated (Fig. 2d,e). The basal ATP production rate in the day 5 IR group is significantly higher compared to untreated (Fig. 2f), which corresponds to a significantly higher mitochondrial ATP production rate (Fig. 2g).

**Figure 2.**
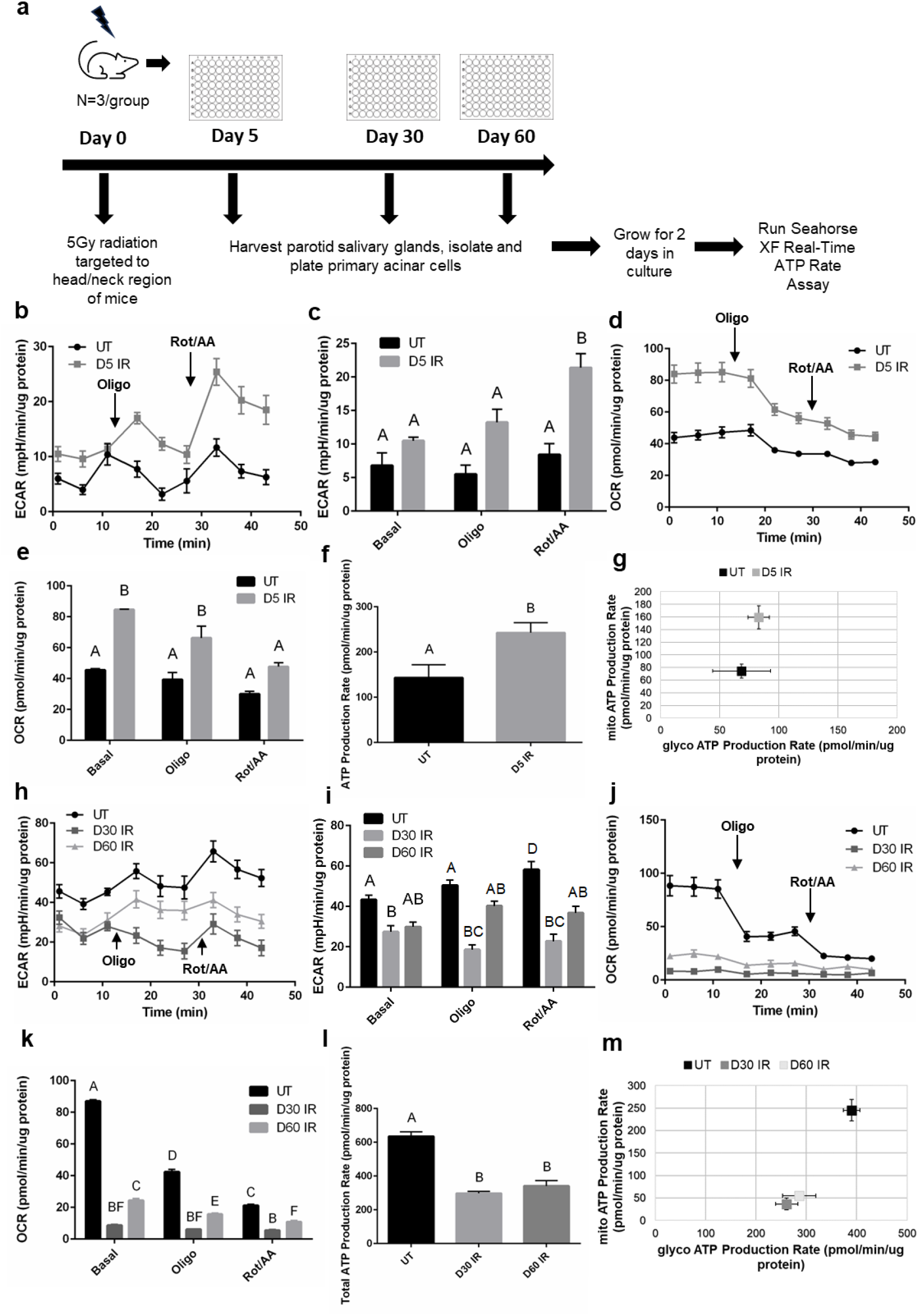
ECAR, OCR, and ATP production increase at 5 days and decrease chronically following radiation in salivary acinar cells. (**a**) Female FVB mice received 5 Gy IR targeted to the head and neck region (n=3/group) and primary acinar cells were isolated from parotid salivary glands of untreated (UT) and IR mice at day 5, 30, and 60 post-IR. Cells were cultured for 2 days prior to running the Seahorse XF Real-Time ATP Rate Assay. (**b**) Individual ECAR readings for untreated UT and day 5 IR (D5 IR) groups with the responses to 2.0uM oligomycin (oligo) injection and 0.5uM rotenone/antimycin A (rot/AA) injection shown. (**c**) Average basal, oligo, and rot/AA ECAR readings for UT and D5 IR. (**d**) Individual OCR readings for UT and D5 IR groups with the responses to 2.0uM oligomycin (oligo) injection and 0.5uM rotenone/antimycin A (rot/AA) injection shown. (**e**) Average basal, oligo, and rot/AA OCR readings for UT and D5 IR. (**f**) Basal ATP production rates of UT and D5 IR groups. (**g**) The proportion of mitochondrial (mito) ATP production rate and glycolytic (glyco) ATP production rate for UT and D5 IR groups. (**h**) Individual ECAR readings for UT, day 30 IR (D30 IR), and day 60 IR (D60 IR) groups with the responses to 2.0uM oligomycin (oligo) injection and 0.5uM rotenone/antimycin A (rot/AA) injection shown. (**i**) Average basal, oligo, and rot/AA ECAR readings for UT, D30 IR, and D60 IR. (**j**) Individual OCR readings for UT, D30 IR, and D60 IR groups with the responses to 2.0uM oligomycin (oligo) injection and 0.5uM rotenone/antimycin A (rot/AA) injection shown. (**k**) Average basal, oligo, and rot/AA OCR readings for UT, D30 IR, and D60 IR. (**l**) Basal ATP production rates of UT, D30 IR, and D60 IR groups. (**m**) The proportion of mitochondrial (mito) ATP production rate and glycolytic (glyco) ATP production rate for UT and D5 IR groups. Each panel is representative of 3 independent assays (n=3/group). Significant differences were determined using one-way ANOVA and Bonferroni post-hoc test, p<0.05. Treatment groups with the same letter are not significantly different from each other.

Cellular bioenergetics change sharply at chronic IR time points. At baseline readings, day 30 and 60 IR ECAR measurements are lower compared to untreated with the day 30 IR group showing a statistically significant decrease (Fig. 2h,i). Following oligomycin and rotenone/antimycin A injections, days 30 and 60 IR ECAR readings continue to remain lower compared to untreated (Fig. 2h,i). At days 30 and 60, baseline OCR measurements are significantly decreased compared to untreated and remain significantly decreased following oligomycin and rotenone/antimycin A injections (Fig. 2j,k). Visual inspection of the cells prior to running the assay and trypan blue staining confirmed viability to ensure that the low OCR cell readings are not due to cell death (Supplemental Figure S2;). Calculation of the basal ATP production rates reveals a significantly lower ATP production rate in the day 30 and day 60 IR groups compared to untreated (Fig. 2l). The day 30 and day 60 IR groups display significantly lower glycolytic and mitochondrial ATP production rates compared to the untreated group (Fig. 2m). Thus, ECAR, OCR, and ATP production rates increase at day 5 IR and then subsequently decrease chronically following radiation.

### Glycolytic enzyme hexokinase protein levels and activity increase in irradiated salivary glands acutely

After identifying changes in cellular ECAR both acutely and chronically in response to radiation treatment, we sought to investigate the key metabolic enzymes that drive glycolysis by measuring protein levels and enzymatic activity of rate-limiting enzymes in salivary gland tissue ^44^. Western blot analysis of the first glycolytic enzyme, hexokinase (HK1-isoform found in the salivary glands), reveals statistically higher protein levels at day 3 IR compared to untreated, day 5 IR, and day 30 IR (Fig. 3a,b). No differences in protein levels of phosphofructokinase (PFKM- isoform found in salivary glands) and pyruvate kinase (PKM1-isoform found in salivary glands) are observed between the untreated and IR groups (Fig. 3a,c-d). The enzymatic activity of hexokinase measured over 60 minutes is higher in the day 3 IR group compared to untreated, day 5 IR, day 14 IR, and day 30 IR groups (Fig. 3e). The enzymatic activity of pyruvate kinase measured over 60 minutes is slightly higher in the day 30 IR group compared to all other groups but is not statistically significant (Fig. 3f). Lactate is produced from pyruvate under anaerobic conditions when pyruvate is not transported into the mitochondria for aerobic respiration ^45^. The lactate concentration of irradiated salivary gland tissue reveals significantly higher levels at days 5 and 14 IR compared to untreated (Fig. 3g).

**Figure 3.**
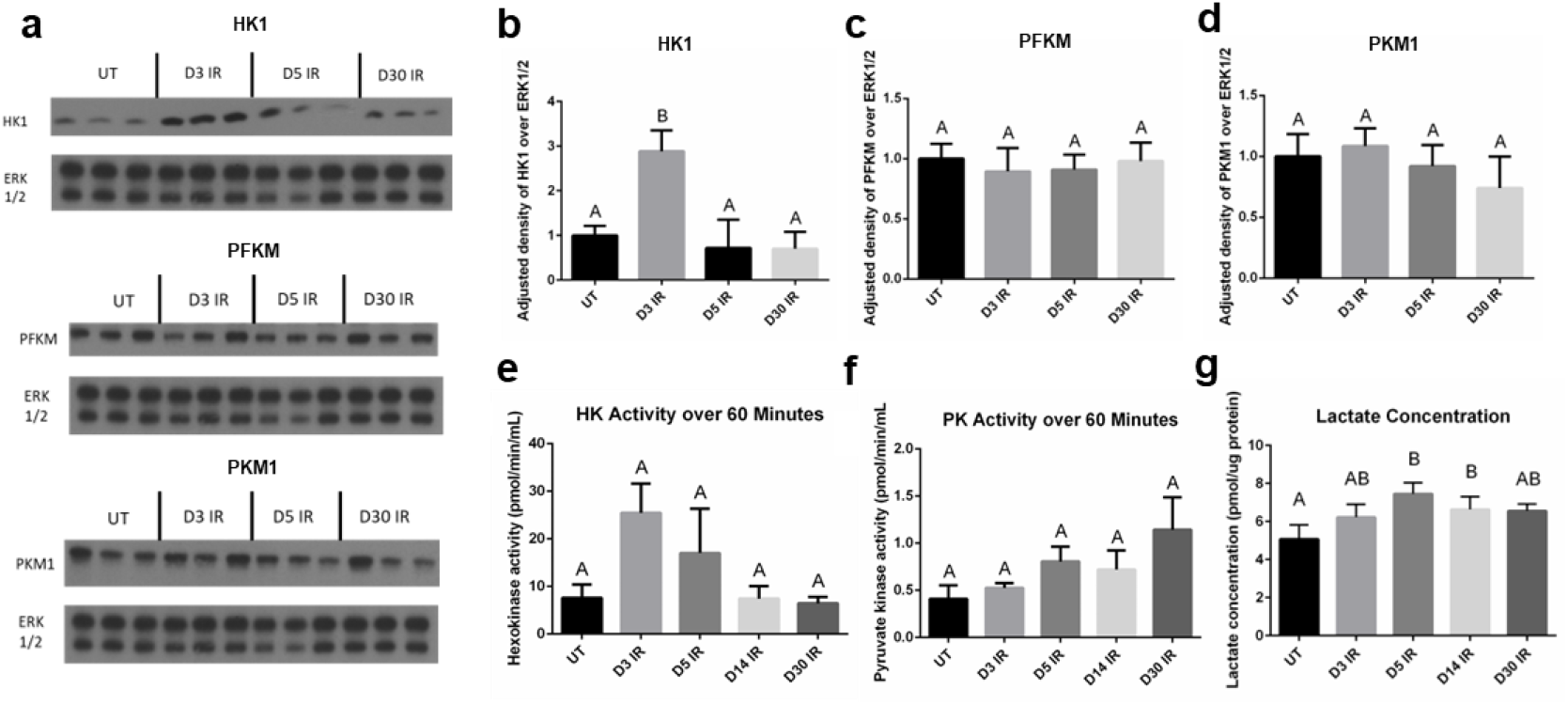
Glycolytic hexokinase protein levels and enzymatic activity increase at day 3 following radiation and lactate concentration increases at day 5 and day 14 following radiation in the salivary gland. Female FVB mice were untreated (UT) or exposed to IR and parotid salivary glands were removed at 3 (D3 IR), 5 (D5 IR), 14 (D14 IR), and 30 (D30 IR) days after radiation treatment (see Fig. 2a). (**a**) Cropped representative western blots probing for hexokinase 1 (HK1), phosphofructokinase-muscle (PFKM), and pyruvate kinase muscle isozyme-1 (PKM1) using untreated, D3 IR, D5 IR, and D30 tissue samples (n=3/group). Extracellular signal-regulated kinase 1/2 (ERK 1/2) was used as a loading control. Original blots shown in Supplementary Figure S5. (**b-d**) Quantification of (a). (**e**) A hexokinase enzyme activity assay was performed on tissue samples (n=4/group) and activity was measured over 60 minutes. (**f**) A pyruvate kinase enzyme activity assay was performed on tissue samples (n=4/group) and activity was measured over 60 minutes. (**g**) A lactate assay kit was used to measure lactate concentration in tissue samples (n=5/group). Data are presented as mean ± standard error of the mean (SEM). Significant differences were determined using one-way ANOVA and Bonferroni post-hoc test, p<0.05. Treatment groups with the same letter are not significantly different from each other.

### Mitochondrial complex I and III protein subunit levels and spare respiratory capacity decrease in day 5 irradiated salivary tissue and acinar cells

Due to the observed changes in OCR at acute and chronic time points in irradiated salivary acinar cells, we further investigated mitochondrial function in salivary gland tissue. In our previous work, we observed significant decreases in transcript families belonging to the core subunit of Complex I in the mitochondrial electron transport chain (*NDUF* and *MT-ND*), and in transcript families belonging to various subunits of Complex III (*UQCR*, *CYC1*, and *MT-CYB*) at day 5 IR in parotid salivary gland tissue ^21^. Therefore, we measured protein levels of Complex I- 75 kDa subunit and of UQCRC2, a subunit of Complex III. We observe a significant decrease in Complex I-75 kDa subunit at day 5 IR compared to untreated (Fig. 4a,b), and we observe a significant decrease in Complex III-UQCRC2 subunit at days 3 and 5 IR (Fig. 4c,d). These data suggest a decrease in mitochondrial respiration at days 3 and 5 IR in parotid salivary gland tissue. To further explore the effects of irradiation on mitochondrial function, we measured spare respiratory capacity in primary salivary acinar cells using the Seahorse XF Cell Mito Stress Test, which has been used to assess mitochondrial dysfunction in a number of disease models ^46,47,48^. Due to the increased OCR and ATP production observed at day 5 IR and the decrease in mitochondrial complex protein I and III subunits, we chose day 5 IR as the time point to investigate spare respiratory capacity. Following FCCP injection, OCR is significantly higher in the untreated group compared to day 5 IR, indicating a higher maximal respiration without radiation treatment (Fig. 4e,f,g). The spare respiratory capacity of the untreated group is significantly higher compared to the day 5 IR group (Fig. 4g), suggesting that day 5 IR salivary acinar cells have a reduced ability to respond to increased energy demand under stress, which is reflective of mitochondrial dysfunction.

**Figure 4.**
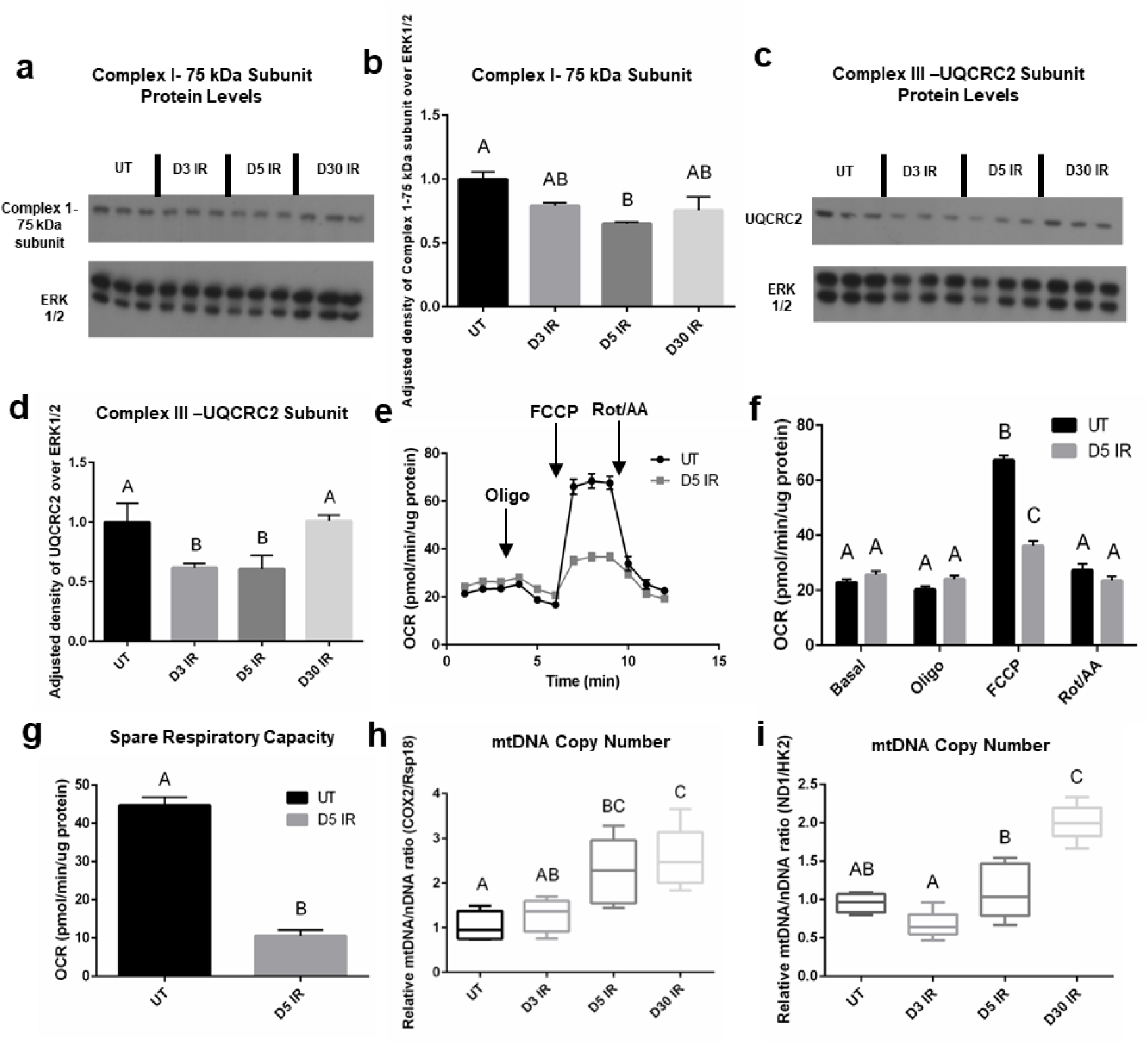
Mitochondrial complex I and III protein subunit levels and spare respiratory capacity decrease at day 5 following radiation and mitochondrial DNA copy number increases chronically following radiation in the salivary gland. Female FVB mice were untreated or exposed to IR and parotid salivary glands were removed at D3, D5, and D30 after radiation (see Fig. 2a). (**a**) Cropped representative western blot probing for complex I-75 kDa subunit using tissue samples (n=3/group). Original blots shown in Supplementary Figure S6. (**b**) Quantification of (a). Data presented as mean ± standard error of the mean (SEM). (**c**) Cropped representative western blot probing for ubiquinol-cytochrome-C reductase complex core protein 2 (UQCRC2) using tissue samples (n=3/group). (**d**) Quantification of (c). Data presented as mean ± standard error of the mean (SEM). (**e-g**) Female FVB mice received 5 Gy IR targeted to the head/neck region (n=3/group) and primary acinar cells were isolated from parotid salivary glands of UT and IR mice at day 5 and cultured for 2 days prior to the Seahorse XF Cell Mito Stress Test. (**e**) Individual OCR readings for UT and D5 IR groups with responses to 2.0uM oligomycin (oligo) injection, 2.0uM FCCP injection, and 0.5uM rotenone/antimycin A (rot/AA) injection shown. (**f**) Average basal, oligo, FCCP, and rot/AA OCR readings for UT and D5 IR. (**g**) Spare respiratory capacity of UT and D5 IR groups. Each panel is representative of 2 independent assays (n=3/group). Significant differences were determined using the two-tailed unpaired T-test, p<0.05 in panel (g). (**h-i**) DNA was isolated from tissue samples (n=5/group) and RT-qPCR was performed with primers specific for the mitochondrial gene cyclooxygenase-2 (*COX2*) and the ribosomal gene ribosomal protein S18 (*Rsp18*) (h), and primers specific for the mitochondrial gene NADH:ubiquinone oxidoreductase core subunit 1 (*ND1*) and the nuclear gene hexokinase 2 (*HK2*) in (i). Data were normalized to 15S ribosomal protein as an internal control and fold change was calculated relative to DNA content in UT mice. Mitochondrial DNA (mtDNA) copy number was estimated using the ratio of *COX2/Rsp18* DNA content (h) and the ratio of *ND1/HK2* DNA content (i). Data presented as interquartile range of the data with the median indicated by the line ± standard error of the mean (SEM). Significant differences were determined using one-way ANOVA and Bonferroni post- hoc test, p<0.05. Treatment groups with the same letter are not significantly different from each other.

### Mitochondrial DNA copy number increases chronically following radiation in the salivary gland

Mitochondrial DNA copy number has been demonstrated to increase following radiation in epithelial and stem cells while oxidative phosphorylation enzyme function decreases, which reflects mitochondrial damage ^49,50,51^. The relative ratio of a mitochondrial DNA target to a nuclear DNA target gives an estimate of mtDNA copy number, and we utilized two different mitochondrial DNA and nuclear DNA targets from studies performed by Zhao et al. and Quiros et al.^52,53^. Looking at the relative ratio of *COX2* mitochondrial gene target to *Rsp18* genomic gene target, we see a significant increase in the relative ratio of *COX2/Rsp 18* at days 5 and 30 IR compared to untreated, with no significant difference observed at day 3 IR (Fig. 4h). Looking at the relative ratio of *ND1* mitochondrial gene target to *HK2* genomic gene target, we observe a significant increase in the relative ratio of *ND1/HK2* at day 30 IR compared to untreated (Fig. 4i). At day 3 IR, the *ND1/HK2* relative ratio appears to slightly decrease compared to untreated, although this decrease is not significant (Fig. 4i). The observed increase in mtDNA copy number suggests mitochondrial dysfunction begins at intermediate time points and continues chronically following irradiation of the salivary gland.

### Dependency on long-chain fatty acids as a fuel source increases at day 5 following radiation in salivary acinar cells

Due to the observed increase in ATP production and reduced spare respiratory capacity at day 5 IR, we investigated the fuel preference of day 5 IR salivary acinar cells using the Seahorse XF Mito Fuel Flex Test Kit ^54^. The fuel dependency of acinar cells to oxidize glucose, glutamine, and long-chain fatty acids was assessed by measuring OCR in response to the following fuel inhibitors: UK5099 inhibits the transport of pyruvate from glycolysis into the mitochondria for oxidation, BPTES inhibits the conversion of glutamine to glutamate so that it cannot enter the Kreb’s cycle for oxidation, and etomoxir inhibits carnitine palmitoyl-transferase 1A (CPT1A), which inhibits the transport of long-chain fatty acids into the mitochondria for oxidation (Fig. 5a). Untreated salivary acinar cells appear highly flexible in their use of fuel for energy production and show a low amount of dependency on glutamine while exhibiting no dependency on glucose and long-chain fatty acids (Fig. 5b,c,d). Radiation treatment does not appear to affect glucose oxidation of primary acinar cells as they remain fully flexible in their use of glucose and do not display dependency on this fuel source (Fig. 5b). Radiation treatment significantly decreases glutamine oxidation and significantly increases flexibility of this fuel source compared to the untreated group (Fig. 5c). In contrast, radiation treatment significantly increases dependency on long-chain fatty acids as evidenced by a 27% dependency of total fuel oxidation compared to 0% dependency in the untreated group. Radiation treatment decreases the flexibility of long-chain fatty acids compared to untreated cells (Fig. 5d). Collectively, day 5 irradiated primary acinar cells demonstrate less dependency on glutamine and more dependency on long-chain fatty acids as a fuel source in comparison to non-irradiated cells (Fig. 5c,d).

**Figure 5.**
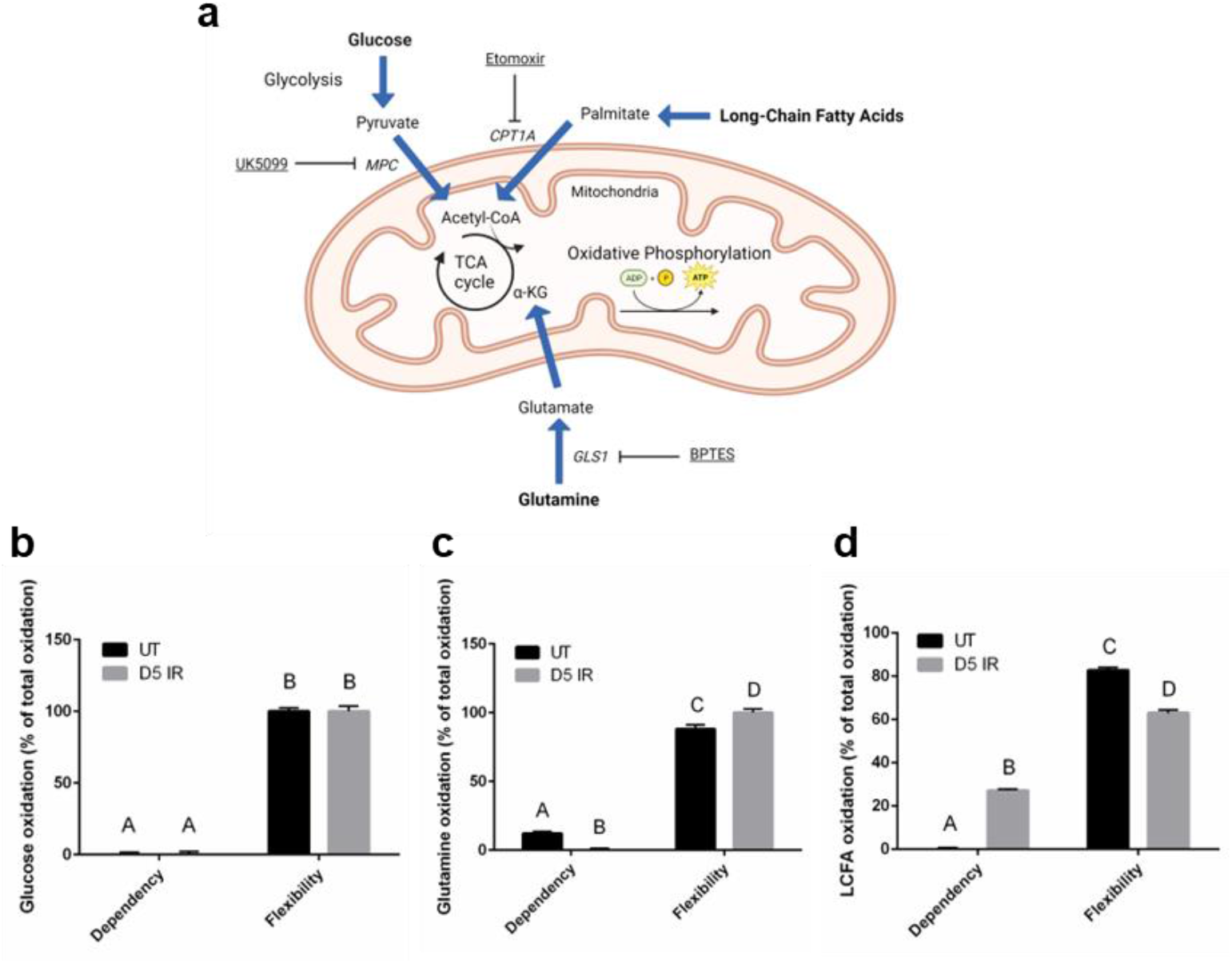
Dependency on long-chain fatty acids as a fuel source increases at day 5 following radiation treatment compared to untreated in salivary acinar cells. Female FVB mice received 5 Gy IR targeted to the head and neck region (n=3/group) and primary acinar cells were isolated from parotid salivary glands of untreated (UT) and IR mice at day 5, cultured for 2 days, and then the Seahorse XF Mito Fuel Flex Test Kit was performed (see Fig. 2a). (**a**) Schematic diagram of glucose, glutamine, and long-chain fatty acid metabolism and where the assay inhibitors block fuel oxidation. Image created in Biorender.com. (**b**) Glucose dependency and flexibility in untreated and day 5 IR (D5 IR) groups. (**c**) Glutamine dependency and flexibility in untreated and D5 IR groups. (**d**) Long-chain fatty acid (LCFA) dependency and flexibility in untreated and D5 IR groups. Data is expressed as % of total oxidation of glucose, glutamine, and long-chain fatty acids ± standard error of the mean (SEM). Significant differences were determined using two-way ANOVA and Bonferroni post-hoc test, p<0.05. Treatment groups with the same letter are not significantly different from each other. Abbreviations: *MPC* = mitochondrial pyruvate carrier complex, *CPT1A* = carnitine palmitoyltransferase 1A, TCA cycle = tricarboxylic acid cycle, α-KG = alpha-ketoglutarate, *GLS1* = glutaminase-1, BPTES = (bis-2-(5-phenylacetamido-1,3,4-thiadiazol-2-yl)ethyl sulfide).

## DISCUSSION

Due to the ongoing need to identify therapeutic targets for restoring salivary gland function following radiation (IR) treatment clinically, mechanistic research is essential for providing new insight. In this study, we investigated the role of energy metabolism and mitochondrial function as mechanistic underpinnings of radiation-induced chronic loss of physiologic function and inefficiencies in wound healing exhibited by parotid salivary glands. Acutely, we observed an increase in glycolysis assessed by extracellular acidification rate (ECAR), oxidative phosphorylation assessed by oxygen consumption rate (OCR), and energy (ATP) production rate at 24 hours post-IR in acinar cells that was attenuated at 48 hours post-IR. At day 5 post-IR when the transition to the chronic damage stage begins, ECAR, OCR, and ATP production rate increased. Energy metabolism reprogramming significantly changed chronically as displayed by decreased ECAR, OCR, and ATP production rate compared to untreated, suggesting changes in energy metabolism underlie the different stages of IR-induced damage in the salivary gland.

It is well established that radiation exposure acutely results in DNA damage and reactive oxygen species (ROS) production ^55,56,57,25,58^, and that mitochondria are the main source of ROS production ^59, 60^. In the salivary gland radiation-damage model, apoptosis peaks at 24 hours post-radiation, and ROS production by mitochondria has been shown to mediate apoptosis ^35,38,61^.

Azimzadeh et al. exposed mice to targeted 3 Gy radiation (IR) to the heart and observed significant upregulation of differentially expressed proteins annotated to mitochondrial oxidative phosphorylation and oxidative stress at 24 hours post-IR ^62^. Another group used a polarographic system to assess mitochondrial oxidative phosphorylation in enterocytes from rats exposed to 1 Gy IR and observed increased oxidative phosphorylation at both 1 and 24 hours post-IR ^63^. These results from *in vivo* radiation models support the observed acute increase in OCR observed in our study and suggest a relationship between increased mitochondrial OCR and ROS production acutely following radiation, which may mediate apoptosis and contribute to tissue dysfunction.

Targeting the acute increase in OCR and ATP production observed in this study may be a mechanism to reduce mitochondrial ROS production and apoptosis and restore secretory function following IR.

The increase in OCR and mitochondrial ATP production observed in day 5 post-IR acinar cells in this study is of interest due to the transition from the acute radiation damage response to the chronic damage response, which is associated with increased compensatory proliferation and mechanistic Target of Rapamycin (mTOR) pathway hyperactivation ^23,64,42^. The protein complex mTORC1 is activated by nutrients, with glutamine, leucine, and arginine classified as the most effective amino acid activators ^65^ . Meeks et al. previously identified significantly increased levels of glutamine in parotid salivary glands at day 5 post-IR ^21^, which may play a role in mTORC1 activation at this time point. mTOR positively regulates gene expression of lactate dehydrogenase, the enzyme that converts pyruvate to lactate ^66^. The significant increase in lactate levels observed at day 5 post-IR in the current study may also contribute to mTORC1 activation. mTOR inhibition has been demonstrated to decrease OCR in cancer cell lines ^67^ , suggesting that mTOR positively regulates OCR. Additionally, mTORC1 activates the translation of nucleus-encoded mitochondrial genes, which further supports mitochondrial respiration ^68^. Our lab previously demonstrated that post-IR treatment with an inhibitor of mTORC1, the rapalogue CCI-779, improves salivary gland function and restores proliferation levels in the glands of mice to untreated levels following IR ^42^. Reprogramming mitochondrial metabolism may be a mechanism by which mTOR inhibition reduces compensatory proliferation and restores salivary gland function chronically following IR. Interestingly, mTOR activates glutaminolysis ^69^, which may increase the flexibility of day 5 irradiated cells to utilize glutamine as a fuel source.

The increased dependency on long-chain fatty acids is likely driving the significant increase in OCR observed at 5 days post-radiation as a higher amount of oxidation-reduction cofactors, flavin adenine dinucleotide (FADH2) and nicotinamide adenine dinucleotide (NADH), are produced from the oxidation of a single long chain fatty acid compared to glucose and glutamine, leading to an increased flux of electrons in the electron transport chain, resulting in increased OCR. ROS are produced from the electron transport chain when electrons leak from complexes I, II, and III and reduce oxygen to superoxide, which can be converted to hydrogen peroxide ^70^ . While the generation of ROS is a normal process that contributes to homeostatic signaling, increases in ROS levels contribute to mitochondrial oxidative stress ^71^. The glutathione antioxidant system is one of the main regulatory systems of hydrogen peroxide levels as glutathione reduces harmful hydrogen peroxide to water and oxygen ^72^ . Decreased glutathione levels are associated with a variety of pathological conditions due to persistently elevated ROS levels ^72^. We previously observed significantly decreased glutathione levels in the parotid salivary gland at 5 days post-radiation ^21^, suggesting that persistently increased ROS levels at this time point are contributing to mitochondrial stress as assessed by decreased spare respiratory capacity in our current study, which would lead to mitochondrial exhaustion chronically ^73^ and contribute to the chronic loss of function phenotype following radiation.

The sharp decrease in OCR observed at chronic IR time points in this study is reflective of mitochondrial oxidative phosphorylation dysfunction. Additionally, the decrease in ECAR and ATP production show an inability to produce sufficient energy to maintain salivary acinar cell function, which may contribute to chronic loss of secretory tissue function. The chronic decrease in energy production and mitochondrial function may be linked to cellular senescence, as dysfunctional mitochondria accumulate in senescent cells due to impaired mitophogy ^74^. Senescent cells become unresponsive to starvation signals and display increased mTOR activity ^75,76^, therefore post-IR treatment with the mTOR inhibitor, the rapalogue CCI-779, may not only restore chronic secretory function by reducing compensatory proliferation^42^ but also by improving mitochondrial dysfunction associated with the senescent phenotype. We observed increases in mtDNA copy number beginning at day 5 and reaching significance at day 30 IR, and these data corroborate Malakhova et al. where they observed an increase in mtDNA copy number in brain and spleen cells of mice exposed to 3 Gy IR ^30^. Malakhova et al. hypothesized that mtDNA repair mechanisms are less efficient compared to nuclear DNA (nDNA) repair mechanisms ^30^, which may be a plausible explanation for why we observe mitochondrial dysfunction occurring chronically in salivary gland tissue. Wu et al. demonstrated that mtDNA stress signaling, displayed by increased mtDNA release, enhances nDNA repair responses ^77^.

Since mtDNA polyploidy is an adaptive repair response ^78^, it may act as a stress signal in response to acute nDNA IR damage in the salivary gland ^57^.

Collectively, this study identified acute changes in energy metabolism following radiation damage that led to chronic energy production impairment and mitochondrial dysfunction in the salivary gland. Limitations to this study include only using a mouse model, which could be improved by using miniature pig or rhesus monkey models to diminish the gap between translation of the animal results to humans. Additionally, we did not include a cumulative radiation damage group in our analysis, which would better mimic the cumulative damage that patients receive during their fractionated radiation treatments. When a patient receives 2 Gy fractionated radiation per day, salivary gland function decreases as evidenced by decreased salivary flow rate (50-60%) within 1 week, reflecting a cumulative total of 10 Gy ^79,80^. The selection of a single 5 Gy dose in a rodent model mimics the low dose acute response when loss of salivary gland function begins clinically in patients ^81,82^. This single 5 Gy dose was also selected since the acute damage response sets up the chronic loss of function response that we are seeking to mechanistically understand ^10^. These findings may be applicable to other exocrine tissues or radiation-damage models for identifying metabolic phenotypes associated with loss of function.

## MATERIALS AND METHODS

### Mice and ethics statement

Female FVB/NJ mice (wild type; stock no. 001800) were purchased from Jackson Laboratories (Bar Harbor, ME). No differences in phenotype have been observed between sexes ^23, 83^. All mice were housed and maintained in accordance with the University of Arizona Institutional Animal Care and Use Committee. All protocols were approved by the Institutional Animal Care and Use Committee (IACUC). The study is reported in accordance with the Animal Research: Reporting In Vivo Experiments (ARRIVE) guidelines.

### Radiation treatment

#### Mice

Four- to eight-week-old female FVB mice were sedated via 20uL intraperitoneal injection of a ketamine-xylazine mixture (70 mg/kg-10 mg/ml) before irradiation. Mice were then constrained in 50-ml conical tubes and shielded with >6 mm lead with only their head and neck region exposed to a single dose of 5 Gy irradiation (X-ray, RS 2000 Small Animal Irradiator, Rad Source). Animals were monitored for 1 hour following radiation treatment to ensure full recovery from the anesthesia.

#### Primary cells

Primary cells isolated from parotid salivary glands were plated on an Agilent Seahorse XF96 Cell Culture Microplate (Part No. 101085-004, Agilent Technologies, Cedar Creek, TX) at a seeding density of 150,000 cells/well and cultured prior to irradiation at 24 hours or 48 hours before running the Seahorse ATP Rate Assay. Cells were exposed to a single dose of 5 Gy irradiation (X-ray, RS 2000 Small Animal Irradiator, Rad Source). The untreated cells were shielded with >6 mm lead.

#### Tissue harvest

Four- to eight-week-old female FVB mice were anesthetized via 50uL intraperitoneal injection of a ketamine-xylazine mixture (70 mg/kg-10 mg/ml). Mice were monitored for 1-5 minutes and when no pedal reflex was observed, the mice were euthanized by exsanguination cutting the renal artery. Parotid salivary glands were immediately harvested from the mice and used for experiments.

#### Primary cell culture

Our primary cell isolation protocol follows the recommendations set by Dr. David O. Quissell who documented the cell culture conditions that select for primary cell acinar enrichment from the salivary glands of rodents in 1986 ^84^.Parotid glands were removed from four-eight-week-old female FVB mice (n=3/group), minced for 2 minutes, and added to a siliconized Erlenmeyer flask with 30 mL dispersion media (Hank’s salt solution, 1 mg/ml collagenase, and 1mg/ml hyaluronidase, pH 7.4). Cells were incubated in a shaking water bath at 37°C for 1 hour with mechanical agitation at 40, 45, 50, 55, and 60 min. Cells were then centrifuged, resuspended in wash media [modified Hank’s solution containing CaCl2 and 0.2% (wt/vol) BSA], recentrifuged, and resuspended again. The suspension was run through a sterile funnel filter, recentrifuged, and suspended in primary cell culture media: DMEM/F12 containing (in wt/vol except where noted) gentamycin (0.5%; Fisher Scientific), fungizone (0.2%; Invitrogen), hydrocortisone (0.04%; Sigma-Aldrich), EGF (0.4%; Fisher Scientific), insulin (0.125%; Invitrogen), transferrin (0.125%, Invitrogen), retinoic acid (0.05%; Sigma-Aldrich), glutamine (1.25%; Invitrogen), nonessential amino acids (1%; Invitrogen), trace elements (1%; Fisher Scientific), and fetal bovine serum (5% vol/vol; Fisher Scientific). The identity of the harvested cells as acinar was confirmed by morphological visual inspection after the harvesting was complete as they display a “cobblestone” structure reflective of epithelial cell cultures ^85^. Cells were plated (150,000 cells/well) on an Agilent Seahorse XF96 Cell Culture Microplate (Part No. 101085-004, Agilent Technologies, Cedar Creek, TX) that was manually coated with 50 ug/mL rat tail collagen I (Ref. no. A10483-01, Gibco, Grand Island, NY) prior to cell plating. The cells sat for 1 hour at room temperature prior to placement in the 37°C/5% CO_2_ incubator for culture. Three mice supply an adequate number of cells for a single treatment group on one Seahorse microplate, with 1-3 treatment groups on a single microplate. One microplate from each independent primary cell culture preparation is considered a replicate for Seahorse assays involving the use of primary cells.

#### Trypan blue staining

Trypan blue staining was performed on untreated and irradiated (D30 IR) primary acinar cells at day 2 in culture to verify the viability of the cells. 0.4% trypan blue was added to the primary salivary acinar cells for 15 seconds and rinsed with PBS once prior to imaging with a Leica DM IL Inverted Phase Contrast Microscope at 10X objective. As a positive control, 30% hydrogen peroxide was added to untreated primary acinar cells at a final concentration of 3% for 24 hours prior to trypan blue staining as previously described.

### Seahorse XF Real-Time ATP Rate Assay

#### Acute IR time points

Primary acinar cells were isolated (see previous methods) from parotid salivary glands of untreated female FVB mice (n=6/plate) and plated on a 96-well collagen-coated Seahorse microplate at a seeding density of 150,000 cells/well. Optimization of seeding density and time in culture was performed prior to experimental runs. The effect of collagen on total protein concentration was determined as this was the normalization method for the extracellular acidification rate (ECAR) and oxygen consumption rate (OCR) data and showed to be non-significant (Supplemental Fig. S3). Cells were cultured for 5 days and irradiated at 24 and 48 hours prior to the assay (see Supplemental Fig. S4 for representative images). Cell culture media was removed and replaced with fresh media every 48 hours. One hour prior to running the assay, cell culture media was replaced with assay media (Seahorse XF DMEM medium pH 7.4, Cat. No. 103575-100, Agilent Technologies, Cedar Creek, TX, supplemented with 1 mM pyruvate, 2 mM glutamine, and 10 mM glucose) and incubated at 37°C. The XF Real-Time ATP Rate Assay Kit (Cat. No. 103592-100, Agilent Technologies, Cedar Creek, TX) was performed according to the manufacturer’s protocol using the Seahorse XF96 Analyzer (Agilent Technologies, Cedar Creek, TX). Optimization of drug concentrations was performed prior to experimental runs. Oligomycin (2uM) was injected into the cell media to inhibit ATP synthase and a mixture of rotenone and antimycin A (0.5uM) was injected into the cell media to inhibit Complex I and III, respectively, of the electron transport chain. Basal ATP production rates were calculated using the Seahorse XF Real-Time ATP Rate Assay Kit Report Generator (Wave Desktop Software, Agilent Technologies, Cedar Creek, TX). Briefly, glycolytic ATP production is calculated from the glycolytic proton efflux rate (PER), which is calculated from the ECAR data, while mitochondrial ATP production is calculated from the OCR data ^34^. Data are presented as ECAR, OCR, and basal ATP production rate normalized to protein concentration. Protein content was determined using the bicinchoninic acid (BCA) assay.

#### Day 5 and chronic IR time points

Female FVB mice received 5 Gy IR targeted to the head and neck region (n=3/group) and primary acinar cells were isolated from parotid salivary glands of UT and IR mice at day 5, day 30, and day 60 post-IR. Cells were seeded on a 96-well collagen-coated Seahorse microplate at a seeding density of 150K/well and cultured for 2 days (optimization of time in culture was determined prior to experimental assays). On day 2 the XF Real-Time ATP Rate Assay Kit (Cat. No. 103592-100, Agilent Technologies, Cedar Creek, TX) was run following the manufacturer’s protocol using the Seahorse XF96 Analyzer as described above.

### Western Blotting

Whole protein lysates from parotid glands of female FVB mice (n=3/group) were harvested and processed for immunoblotting as previously described ^23, 86^. The Coomassie Plus-The Better Bradford Assay (Thermo) was used to determine protein concentrations and 50ug total lysate was loaded onto a 10% polyacrylamide gel, transferred to an Immobilon PDVF membrane (Millipore, Bedford, MA), and blocked in 5% nonfat milk in Tris-buffered saline-Tween 20 (1X TBST). The following antibodies were used: anti-hexokinase isoform 1 (HK1) (Product no. C35C4, Cell Signaling), anti-muscle phosphofructokinase (PFKM) (Cat. No. 55028-1-AP, Proteintech), anti-muscle pyruvate kinase isoform 1 (PKM1) (Product no. D30G6, Cell Signaling), anti-complex I – 75 kDa subunit (ABN302, EMD Millipore), anti-complex III ubiquinol-cytochrome C reductase core protein 2 (UQCRC2) subunit (Cat. No. 14742-1-AP, Proteintech), and anti-extracellular signal-regulated protein kinases 1/2 (ERK1/2) (p44/42 MAPK, Cell Signaling). For detection, ECL substrate (Thermo Scientific) or SuperSignal West Pico Chemiluminescent Substrate (Thermo Scientific) was used. Restore Western Blotting Stripping Buffer (Fisher) was used to strip the membrane, re-block with 5% nonfat milk in 1X TBST, and re-probe for the loading control ERK1/2. Densitometry was performed using ImageJ software (NIH).

### Enzyme Activity Assays

A split-sample approach was used for the enzyme activity assays. From the same mouse sample (n=4/group), one parotid salivary gland was analyzed using the hexokinase assay kit (ab136957, Abcam, Cambridge, UK) and the second parotid salivary gland was analyzed using the pyruvate kinase activity assay kit (cat. No. MAK072, Sigma-Aldrich, St. Louis, MO). For both activity assays, the samples were plated on a 96-well plate at a concentration of 10 ng/uL/well.

Luminescence readings were measured every 5 minutes for a total period of 60 minutes to calculate enzymatic activity following the manufacturer’s protocols.

### Lactate Assay

A pair of parotid salivary glands were harvested from each mouse (n=5/group) and directly placed in 500 uL of Lactate Assay Buffer from the L-Lactate Assay Kit (ab65331, Abcam, Cambridge, UK), 10 uL of protease inhibitor (PI) cocktail for mammalian cells (cat # P8340, Sigma-Aldrich, St. Louis, MO) and 5 uL of 100 mM phenylmethylsulfonyl fluoride (PMSF). Samples were homogenized and deproteinated prior to the assay following the manufacturer’s protocol. For the assay, 30 uL of sample was added to a well for a single replicate. Lactate concentration was measured following the manufacturer’s protocol (ab65331, Abcam, Cambridge, UK).

### Quantitative real-time polymerase chain reaction (qRT-PCR)

Parotid glands were removed from mice (n=5/group) and total DNA was immediately isolated using the DNeasy Blood & Tissue Kit (Ref. No. 69504, Qiagen, Hilden, Germany). DNA samples were diluted with nuclease-free water to a concentration of 10 ng/uL. For quantitative (q) PCR, samples were analyzed in triplicate for each DNA sample (five mice per condition) with an iQ5 Real-Time PCR Detection System (Bio-Rad). Master mixes were prepared as follows: 5 μl of diluted DNA, 2 μl of the forward and reverse primers, 7.5 μl of SYBR Green (Qiagen), and 0.5 μl nuclease-free water to a final volume of 15 μl. Forty cycles of PCR were performed (95°C for 15 s, 54°C for 30 s, 72°C for 30 s); fluorescence detection occurred during the 72°C step at each cycle. The data were analyzed using the 2-ΔΔCT method ^87^. Results were normalized to S15, which remains unchanged in response to treatment, and graphed as fold change in comparison to untreated. The following primers were purchased from Integrated DNA Technologies (Coralville, IA): *S15* (ribosomal protein S15; Forward: 5′-ACT ATT CTG CCC GAG ATG GTG-3′; Reverse: 5′-TGC TTT ACG GGC TTG TAG GTG-3′), *COX2* (cyclooxygenase-2; Forward: 5′-ATA ACC GAG TCG TTC TGC CAA T-3′; Reverse: 5′-TTT CAG AGC ATT GGC CAT AGA A-3′), *Rsp18* (ribosomal protein S18; Forward: 5′-TGT GTT AGG GGA CTG GTG GAC A-3′; Reverse: 5′-CAT CAC CCA CTT ACC CCC AAA A-3′), *HK2* (hexokinase 2; Forward: 5′-GCC AGC CTC TCC TGA TTT TAG TGT-3′; Reverse: 5′- GGG AAC ACA AAA GAC CTC TTC TGG-3′), and *ND1* (NADH:Ubiquinone Oxidoreductase Core Subunit 1; Forward: 5′-CTA GCA GAA ACA AAC CGG GC-3′; Reverse: 5′-CCG GCT GCG TAT TCT ACG TT-3′.

### Mitochondrial DNA (mtDNA) Copy Number

Total DNA was extracted from mouse parotid gland tissue (n=5/group) following the manufacturer’s protocol for DNeasy Blood & Tissue Kit (Ref. No. 69504, Qiagen, Hilden, Germany). Mitochondrial DNA (mtDNA) copy number was measured by assessing the relative levels of mitochondrial gene targets *COX2* and *ND1* to genomic DNA (gDNA) targets *Rsp18* and *HK2* ^52, 53^ in extracts of total DNA using qRT-PCR analysis as described above.

### Seahorse XF Mito Fuel Flex Test and Cell Mito Stress Test

Female FVB mice (n=3/group) received 5 Gy IR targeted to the head and neck region and primary acinar cells were isolated from parotid salivary glands of UT and IR mice at day 5 post-IR. Cells were seeded on a 96-well collagen-coated Seahorse plate at a seeding density of 150,000 cells/well and cultured for 2 days. One hour prior to running the assay, cell culture media was replaced with assay media (Seahorse XF DMEM medium pH 7.4, Cat. No. 103575-100, Agilent Technologies, Cedar Creek, TX, supplemented with 1 mM pyruvate, 2 mM glutamine, and 10 mM glucose) and incubated at 37°C.

### Mito Fuel Flex Test

The Seahorse XF Mito Fuel Flex Test Kit (Cat. No. 103260-100, Agilent Technologies, Cedar Creek, TX) was performed according to the manufacturer’s protocol using the Seahorse XF96 Analyzer (Agilent Technologies, Cedar Creek, TX). UK5099 (2 uM) was injected into the cell media to inhibit glucose oxidation, BPTES (3 uM) was injected into the cell media to inhibit glutamine oxidation, and Etomoxir (4 uM) was injected into the cell media to inhibit long-chain fatty acid oxidation. The order of inhibitor injections and inhibitor combinations allowed the calculation of fuel dependency (glucose, glutamine, and long-chain fatty acids) and the calculation of the capability of the cells to use alternate fuel sources. Fuel flexibility was calculated using the Seahorse XF Mito Fuel Flex Test Report Generator (Wave Desktop Software, Agilent Technologies, Cedar Creek, TX) and data is presented as oxygen consumption rate (OCR) normalized to protein concentration. Protein content was determined using the BCA assay.

### Cell Mito Stress Test

The Seahorse XF Cell Mito Stress Test Kit (Cat. No. 103015-100, Agilent Technologies, Cedar Creek, TX was performed according to the manufacturer’s protocol using the Seahorse XF96 Analyzer (Agilent Technologies, Cedar Creek, TX). FCCP concentration was optimized prior to the experimental runs. Oligomycin (2uM) was injected into the cell media to inhibit ATP synthase, carbonyl cyanide-4 (trifluoromethoxy) phenylhydrazone (FCCP) (2uM) was injected into the cell media to uncouple electron transport from ATP production in the electron transport chain, and a mixture of rotenone and antimycin A (0.5uM) was injected into the cell media to inhibit Complex I and III, respectively, of the electron transport chain. Spare respiratory capacity was calculated using the Seahorse XF Cell Mito Stress Test Report

Generator (Wave Desktop Software, Agilent Technologies, Cedar Creek, TX) and data is presented as oxygen consumption rate (OCR) normalized to protein concentration. Protein content was determined using the BCA assay.

### Statistics

Data were analyzed using Prism 6.04 (GraphPad, La Jolla, CA). All values are reported as means ± standard error (SE) of at least three independent experiments unless denoted otherwise. Statistical tests were two-sided and differences between more than two group means were evaluated using the one-way analysis of variance (ANOVA) test and Bonferroni post-hoc multiple comparison test with significant differences at *p* < 0.05. Differences between two group means were evaluated using two-sided unpaired T-tests with significant differences at *p* < 0.05. Treatment groups with the same letter are not statistically different from each other.

## Supplemental Data

All supplemental data can be accessed using the link provided by the journal.

## Data Availability

The datasets generated and analyzed during the current study are available in the Zenodo repository using the following link: https://zenodo.org/doi/10.5281/zenodo.8157794.

## ACKNOWLEDGEMENTS

We acknowledge members of the Kirsten Limesand and Sean Limesand labs as well as Anup Srivastava, Jessica Martinez, Floyd “Ski” Chilton, and Ningning Zhao for useful discussions. We thank the Schnellmann lab for training and allowing access to their Seahorse XF96 Analyzer. This work was supported in part by NIH DE023534 and DE029166 to Kirsten Limesand and stipend support for Lauren Buss was provided by NIH DE030663-01.

## AUTHOR CONTRIBUTIONS

K.H.L. and L.G.B. designed the study. L.G.B. wrote the manuscript. L.G.B. and B.A.R. collected and analyzed the data. L.G.B. generated the figures. L.G.B. and K.H.L. reviewed the manuscript. K.H.L. edited the manuscript in preparation for its final version.

