## Supplemental Figures for "Radiation-Induced Changes in Energy Metabolism Result in Mitochondrial Dysfunction in Salivary Glands"

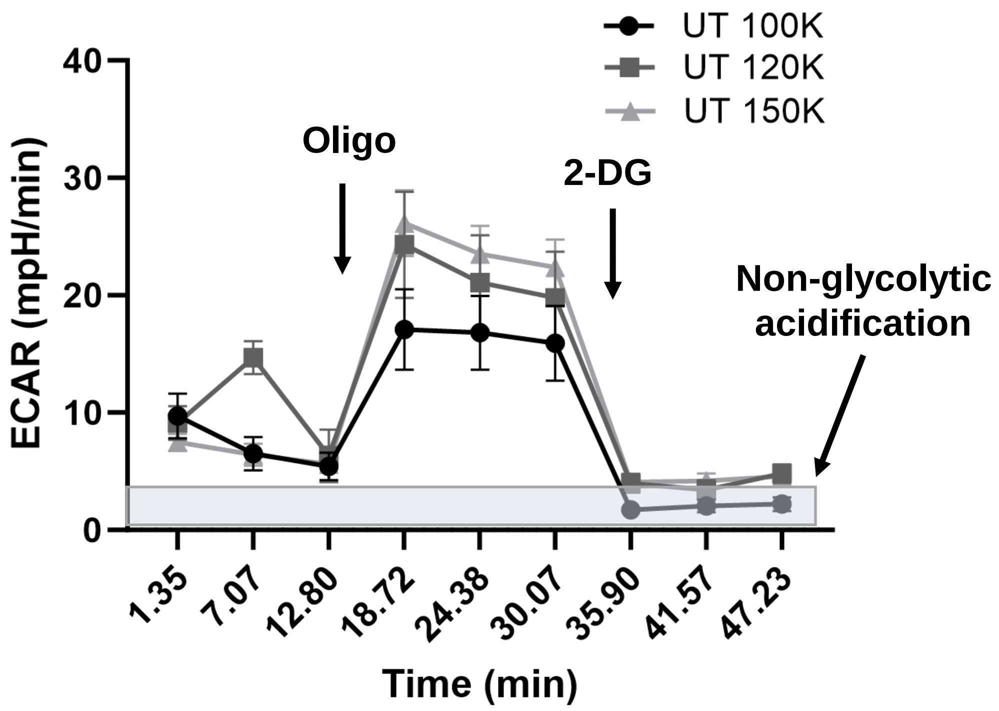


**Figure S1. Seahorse XF optimization experiment for seeding density (100K, 120K, and 150K) of untreated (UT) primary acinar cells on collagen-coated Seahorse XF96 microplates shows minimal non-glycolytic acidification contributing to ECAR**. ECAR measurements following oligomycin injection (2uM) reflect glycolytic capacity as oligomycin inhibits ATP synthase, resulting in a compensatory increase in glycolysis. 2-deoxyglucose (2-DG) is a competitive inhibitor of glucose and following 2-DG injection (50mM), ECAR correspondingly decreases. The difference between ECAR measurements following 2-DG injection and 0 reflect the contribution of non-glycolytic acidification to ECAR measurements, which is less than 5 mpH/min across all seeding densities.

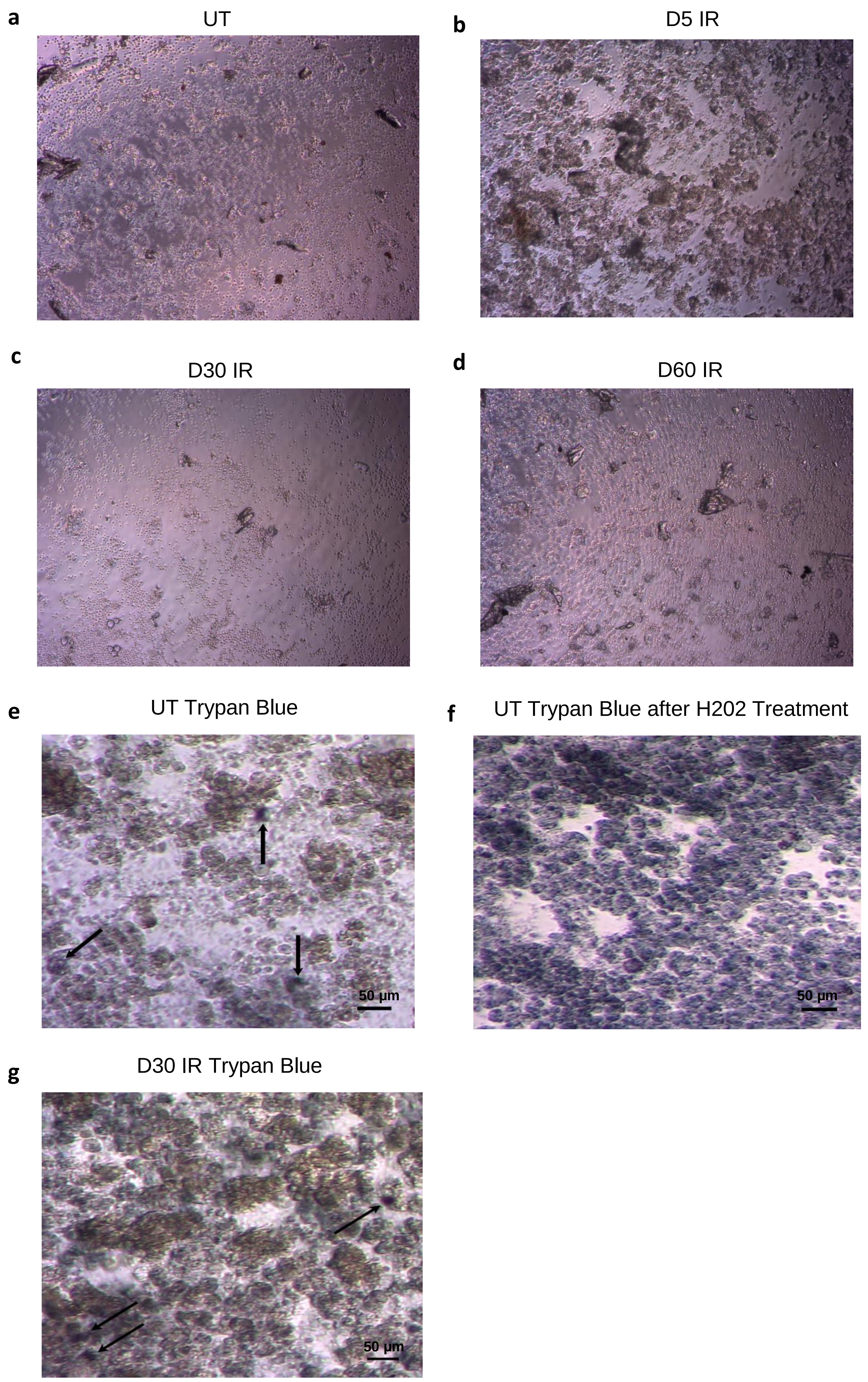


**Figure S2. Representative images of primary salivary gland acinar cells at 2 days in culture viewed using a Leica DM IL Inverted Phase Contrast Microscope at 10X objective. (a)** Untreated (UT) salivary acinar cells. **(b)** Day 5 irradiated (D5 IR) salivary gland acinar cells. **(c)** Day 30 irradiated (D30 IR) salivary gland acinar cells. **(d)** Day 60 irradiated (D60 IR) salivary gland acinar cells. **(e)** 0.4% trypan blue was added to UT salivary acinar cells for 15 seconds and rinsed with PBS once prior to imaging. **(f)** 30% hydrogen peroxide (H202) was added to the untreated salivary acinar cells at a final concentration of 3% for 24 hours prior to trypan blue staining as performed in (e). **(g)** 0.4% trypan blue was added to D30 IR salivary acinar cells for 15 second and rinsed with PBS once prior to imaging. Arrows point to trypan blue-positive cells.

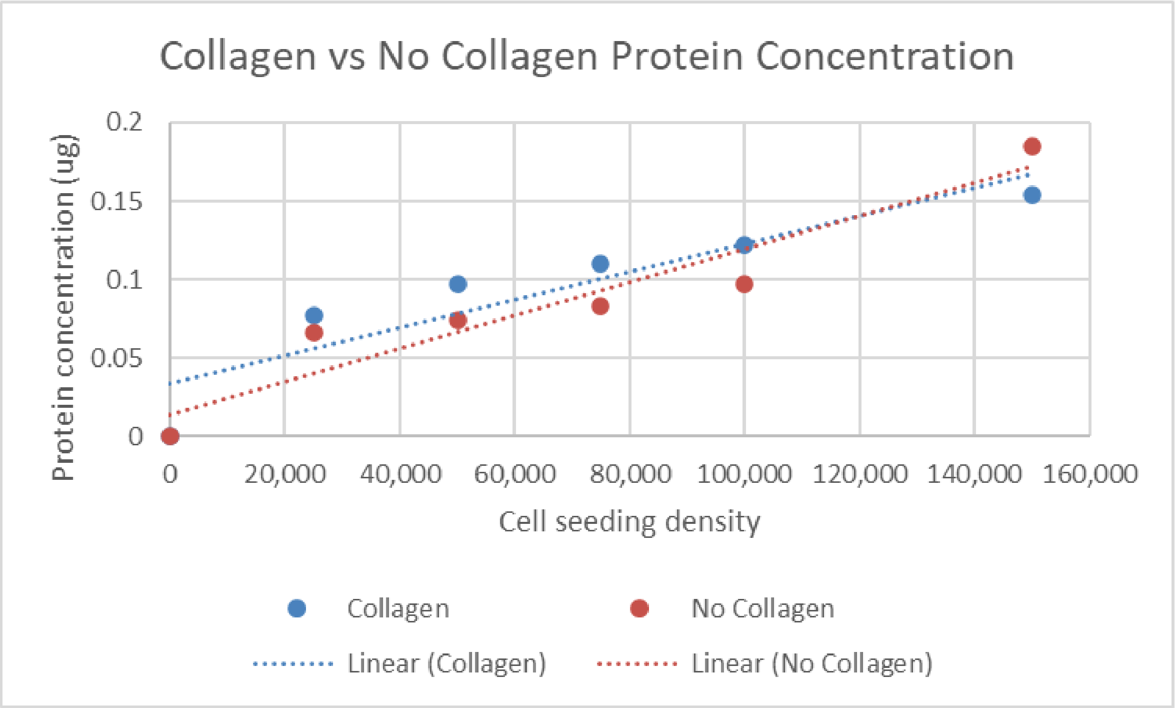


**Figure S3. Similar positive correlations are observed between cell seeding density and protein concentration (ug) for primary acinar cells on Seahorse XF microplates with collagen coating compared to primary acinar cells on Seahorse XF microplates without collagen coating.** The Bicinchoninic acid (BCA) assay was performed on primary acinar cells at 5 days in culture on a Seahorse XF96 microplate coated with 50ug/mL of collagen and on primary acinar cells at 5 days in culture on a Seahorse XF96 microplate without collagen. The correlation coefficient of the linear equation for cell seeding density against protein concentration on collagen coated microplates is $R^2 = 0.841$ and the correlation coefficient of the linear equation for cell seeding density against protein concentration on microplates without collagen is $R^2 = 0.903$.

a UT

b 24hr IR

c 48hr IR

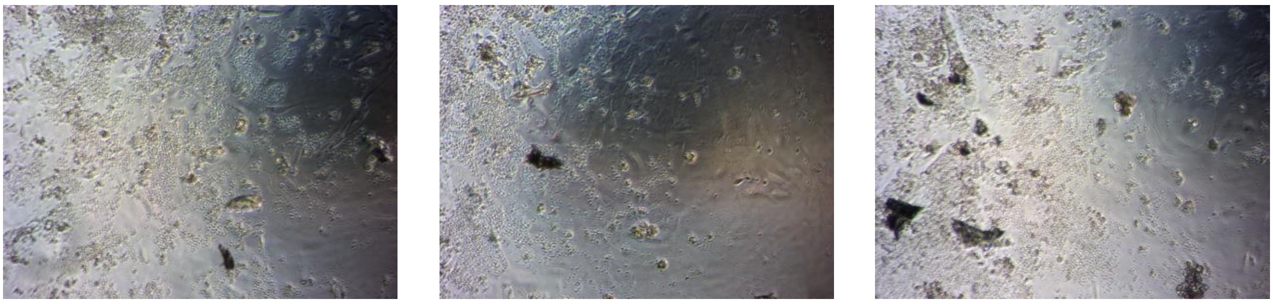

**Figure S4. Representative images of primary salivary gland acinar cells at 5 days in culture viewed using a Leica DM IL Inverted Phase Contrast Microscope at 10X objective. (a)** Untreated (UT) salivary acinar cells. **(b)** 24-hour irradiated (24hr IR) salivary gland acinar cells. **(c)** 48-hour irradiated (48hr IR) salivary gland acinar cells.

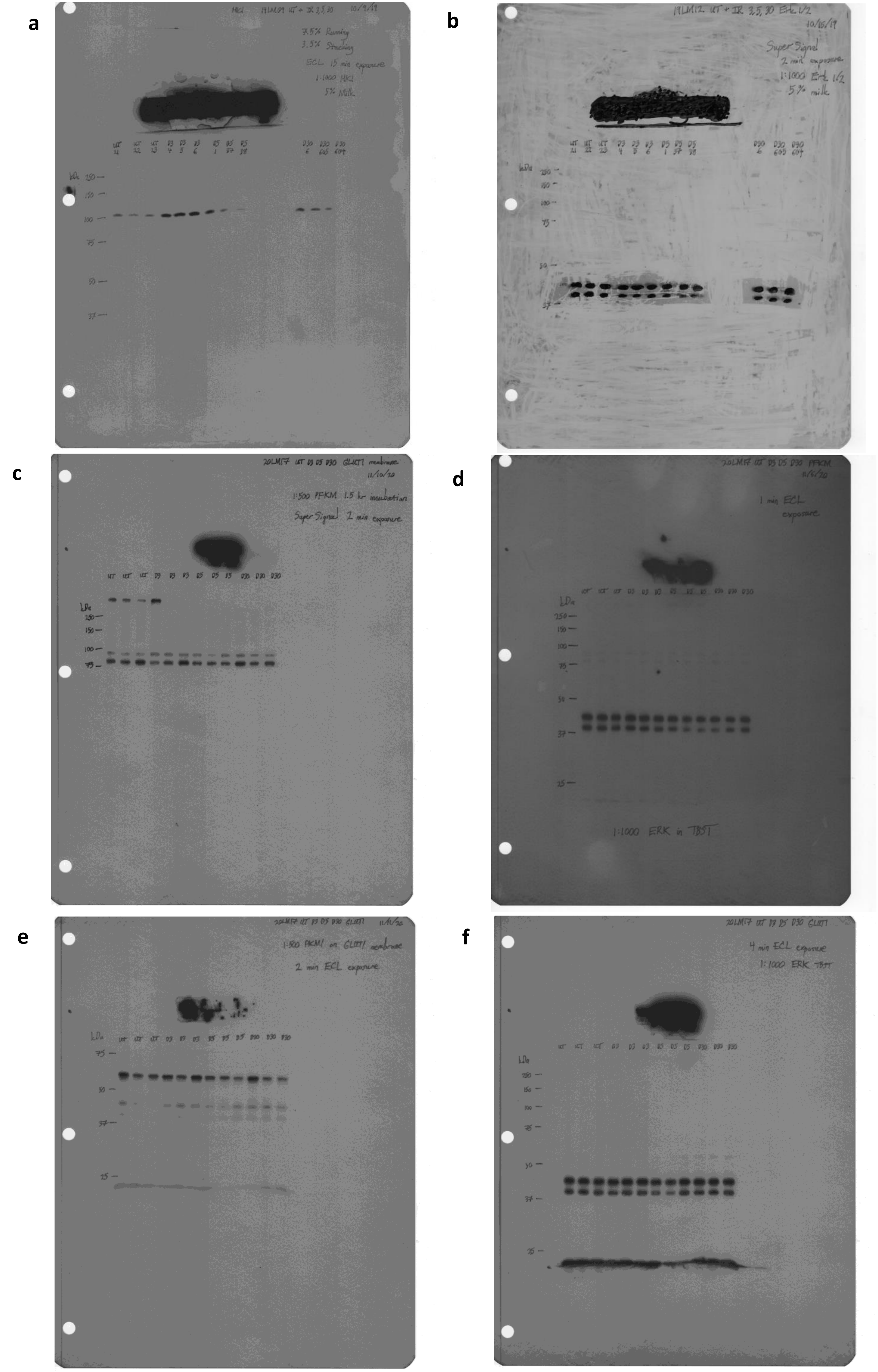


**Figure S5. Original representative western blots from Fig. 3 using untreated (UT), day 3 (D3), day 5 (D5), and day 30 (D30) mouse parotid tissue samples (n=3/group). (a)** Probing for hexokinase 1 (HK1). **(b)** Probing for ERK1/2 on the same blot as in (a). **(c)** Probing for phosphofructokinase-muscle (PFKM). **(d)** Probing for ERK1/2 on the same blot as in (c). **(e)** Probing for pyruvate kinase muscle isozyme-1 (PKM1). **(f)** Probing for ERK1/2 on the same blot as in (e). Membranes were exposed to autoradiography film (Genesee, no. 30-810) and developed using an Srx-101A X-ray film processor (Konica). Membranes were stripped using Restore Western Blotting Stripping Buffer (Fisher, no. 21063), blocked in 5% nonfat milk, and re-probed for loading controls. Densitometry was performed using ImageJ software (NIH).

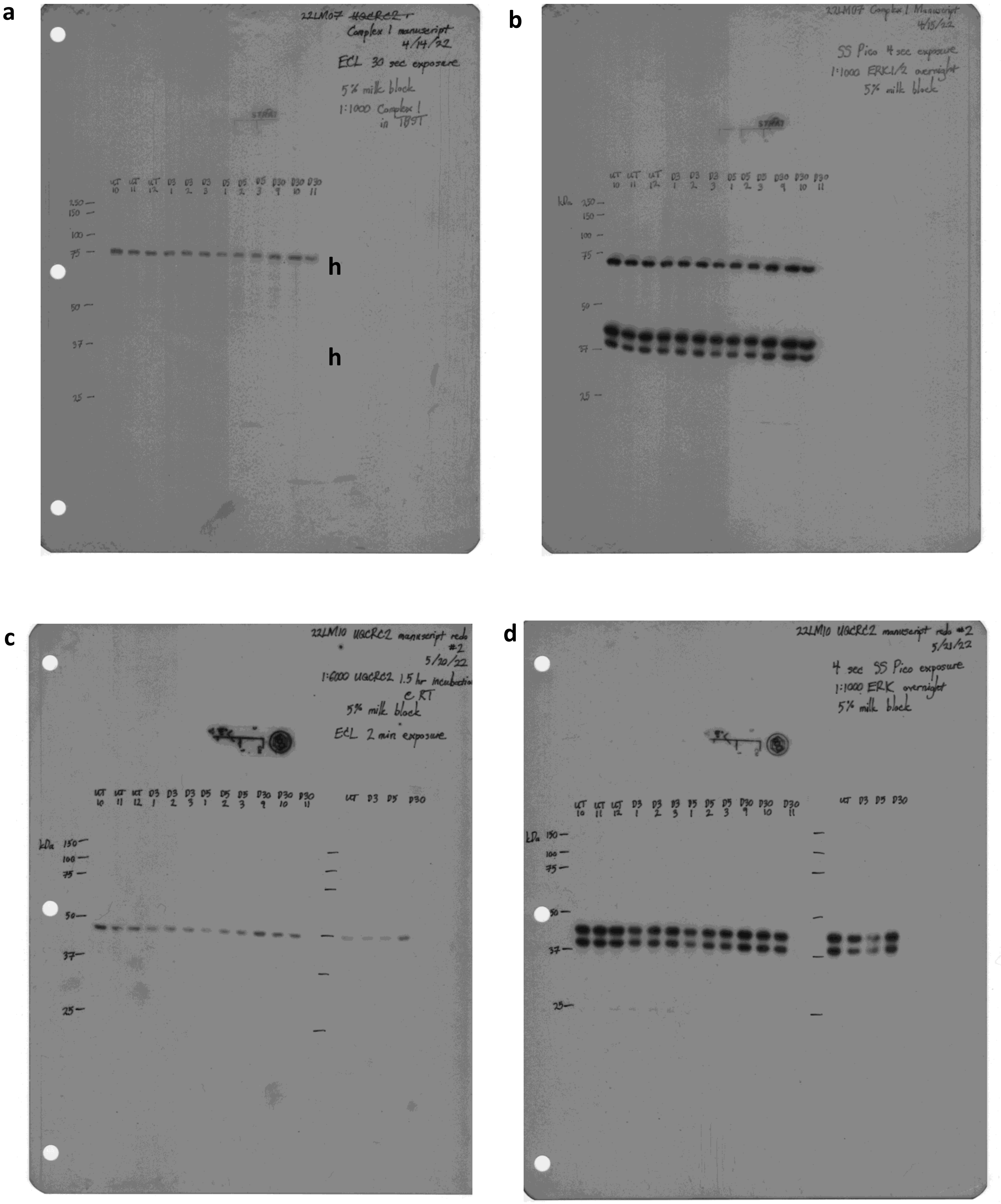


**Figure S6. Original representative western blots from Fig. 4 using untreated (UT), day 3 (D3), day 5 (D5), and day 30 (D30) mouse parotid tissue samples (n=3/group). (a)** Probing for complex I-75 kDa subunit. **(b**) Probing for ERK1/2 on the same blot as in (a). **(c)** Probing for ubiquinol-cytochrome-C reductase complex core protein 2 (UQCRC2). **(d)** Probing for ERK1/2 on the same blot as in (c). Membranes were exposed to autoradiography film (Genesee, no. 30-810) and developed using an Srx-101A X-ray film processor (Konica). Membranes were stripped using Restore Western Blotting Stripping Buffer (Fisher, no. 21063), blocked in 5% nonfat milk, and re-probed for loading controls. Densitometry was performed using ImageJ software (NIH).
